# Heterozygous Lepr Mutation Accelerates Alzheimer-like Neurodegeneration in db/+ Mice via CDK5 Hyperactivation

**DOI:** 10.1101/2025.08.28.672812

**Authors:** Sangita Paul, Juhi Bhardwaj, Srishti Sharma, KT Mohammed Swalih, Deepanshi Koul, B.K. Binukumar

## Abstract

Leptin signaling has neuroprotective effects and is increasingly linked to Alzheimer’s disease (AD). Beyond metabolism, leptin modulates β-amyloid metabolism, tau phosphorylation, and synaptic plasticity. While homozygous Lepr mutations are well studied, the impact of heterozygous mutations on neurodegeneration is unclear. To assess partial Lepr loss, one-year-old db/+ mice were evaluated for metabolic, behavioral, and neuropathological changes. Tests included glucose tolerance, memory assays, Aβ_42_ and tau levels, CDK5 activity, and transcriptomics. Human LEPR variants were curated and classified using ACMG guidelines. Aged db/+ mice showed metabolic dysfunction, cognitive deficits, and AD-like pathology. Compared to controls, db/+ mice had increased body weight, insulin resistance, memory impairments, elevated Aβ_42_, tau hyperphosphorylation, CDK5 hyperactivation, and astrocyte activation. Transcriptomics revealed altered synaptic and mitochondrial pathways. Thirty-three pathogenic or likely pathogenic human LEPR variants were identified. Lepr haploinsufficiency contributes to age-related cognitive decline and AD-like pathology, suggesting it as a genetic risk factor and therapeutic target.

## 1. INTRODUCTION

As per the World Health Organisation (WHO), more than 1.9 billion adults worldwide are overweight, and of these, over 650 million are obese-more than the entire population of Russia and Australia put together. The excess weight acts as a hotbed for a multitude of health problems, like Type 2 diabetes mellitus (T2DM), cardiovascular disease ^1^, certain cancers, obstructive sleep apnea (OSA), respiratory ailments, reproductive issues, osteoarthritis, and mental health issues. The most understudied of them all is cognitive impairment caused by dysregulation of glucose metabolism in obese individuals. Studies increasingly reveal a strong association between a higher mid-life body mass index (BMI), impaired brain function, and cognitive deterioration ^2^ ^3^.

To better understand the underlying mechanisms of diabetes-induced cognitive impairment and to test new therapy approaches, a suitable animal model of the condition is imperative. The obese and insulin-resistant db/db mice, which have a leptin-receptor deficit, are one of the most common laboratory animals used for obesity and diabetes research. Because of a mutation in the leptin receptor gene (*Lepr*), db/db mice are a good genetic model for T2DM. In homozygous (db/db) mice, the mutation is believed to be autosomal recessive with complete penetrance that results in metabolic abnormalities that mimic diabetic mellitus in humans. Db/db mice also replicate neurological pathology, as seen in T2DM patients ^4^. Studies have shown that db/db mice exhibit cognitive impairment in various tasks related to learning and memory. For example, they show deficits in spatial memory, object recognition memory, and fear conditioning memory ^5^. Db/db mice also show hallmarks of neurodegeneration, like elevated amyloid beta and tau hyperphosphorylation ^5,6^. CDK5, an atypical kinase that is dysregulated in neurodegenerative disorders, is also found to be dysregulated in db/db mice ^4^.

The heterozygous (db/+) mice are often used as controls for db/db mice studies. However, research conducted on db/+ mice indicates that there is a decrease in leptin signalling with age, which can be due to a lower quantity of intact long receptor isoform molecules ^7^. Also, it was reported that Db/+ mice develop very high plasma leptin levels in old age compared to wild-type mice ^8–10^. The implication is that the *Lepr* gene is not entirely recessive, and heterozygosity at the leptin receptor may contribute to vulnerability to environmental factors that promote obesity and, possibly, cognitive impairment ^11^.

To the best of our knowledge, no comprehensive study has examined the potential impact of the db/+ mutation on the cognitive abilities of aged db/+ mice in comparison to age-matched wild type control mice. Hence, the present study focussed on exploring the impact of heterozygous db/+ mutation on cognitive abilities and pathological changes in the mice brain. To extend the relevance of our findings to the human population, we systematically compiled and classified all reported human *LEPR* variants according to the American College of Medical Genetics and Genomics (ACMG) guidelines. This comprehensive analysis allowed us to determine the allele frequencies of pathogenic and likely pathogenic variants in the population.

## 2. RESULTS

### 4.1 *Lepr* haploinsufficiency increases susceptibility to obesity and type 2 diabetes in aged mice

This study investigated the impact of a heterozygous Lepr mutation on obesity, glucose regulation, and cognitive function in aged db/+ mice. We confirmed the genotypes of db/+ and control mice using restriction site mapping and Sanger sequencing. Notably, the G to T mutation in the Lepr gene responsible for introducing a new RsaI restriction site in db/db mice was validated in db/+ mice (see Figure 1C) and further confirmed by Sanger sequencing (Figure 1D).

**Figure 1.**
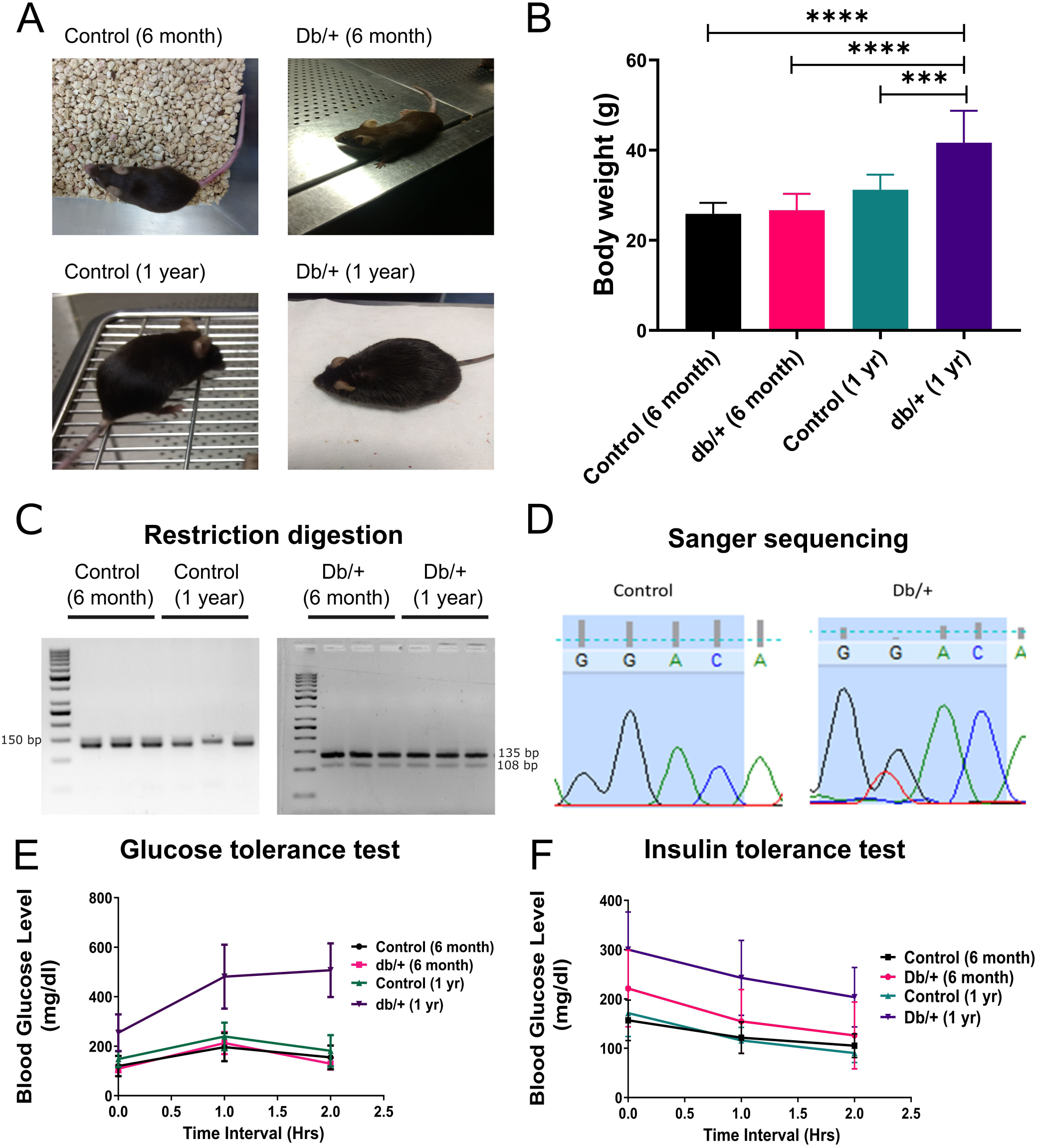
Heterozygous Lepr mutation in aged db/+ mice leads to obesity and high blood glucose. The generation of db/+ models and their validation (A) Representative images of control groups six months, one year, db/+ six months, and db/+ one year. (B) Bar graph representing the body weights of all the mouse groups. Genotyping of the control and heterozygous db/+ mice through (C) Restriction digestion and (D) Sanger sequencing of the mutated site GGAC. Biochemical parameters showing (E) GTT (F) ITT of the control vs db/+ mice group of 6 months, as well as 1 year. All the groups have n=9 mice, and the data was presented as mean ± SD. The analysis was done using one-way ANOVA, and significance was denoted as *p>0.05, **p>0.01; ***p>0.001

Metabolic assessments were then conducted using glucose tolerance tests (GTT) and insulin tolerance tests (ITT). At six months of age, the body weights of db/+ mice were similar to those of controls. However, by one year, db/+ mice exhibited a significantly higher body weight compared to controls (Figures 1A and 1B). These results suggest that although the db mutation is present from birth, its impact on weight gain becomes more evident with age. Initially, compensatory mechanisms such as efficient energy expenditure or effective satiety signaling may mitigate the mutation’s impact, but these protective processes may deteriorate over time, leading to increased adiposity.

In terms of glucose regulation, no significant differences were observed between control and db/+ mice at six months. Yet, in aged db/+ mice, glucose tolerance was impaired, as shown by elevated blood glucose levels at the 2-hour mark and an increased area under the curve during the GTT (Figure 1E). Moreover, the ITT revealed that db/+ mice had consistently higher insulin levels at all time points measured (0, 0.5, 1, 1.5, 2.0, and 2.5 hours) compared to controls, suggesting a decrease in insulin sensitivity (Figure 1F).

Overall, these findings highlight that the heterozygous Lepr mutation plays a significant role in the age-dependent development of obesity and type 2 diabetes, emphasizing the importance of longitudinal studies in understanding genetic influences on metabolic health.

### 4.2 Lepr haploinsufficiency Leads to Reduced Muscle Strength, Impaired Coordination, and Increased Anxiety in Aged Mice

Given the metabolic disturbances observed in aged db/+ mice carrying the heterozygous *Lepr* mutation, we extended our investigation to explore how this mutation affects general behavior, motor coordination, muscle strength, and anxiety-like behaviors across different age groups— specifically at six months and one year. To assess sensorimotor coordination and motor learning, we employed the rotarod test. While no significant differences were detected between db/+ and wild-type mice at six months, one-year-old db/+ mice exhibited a marked decline in both sensorimotor coordination and muscle strength compared to their age-matched wild-type counterparts (Figure 2 A-B). This impairment was further supported by the grip strength test, which revealed a significant reduction in forelimb strength in older db/+ mice (Figure 2C).

**Figure 2.**
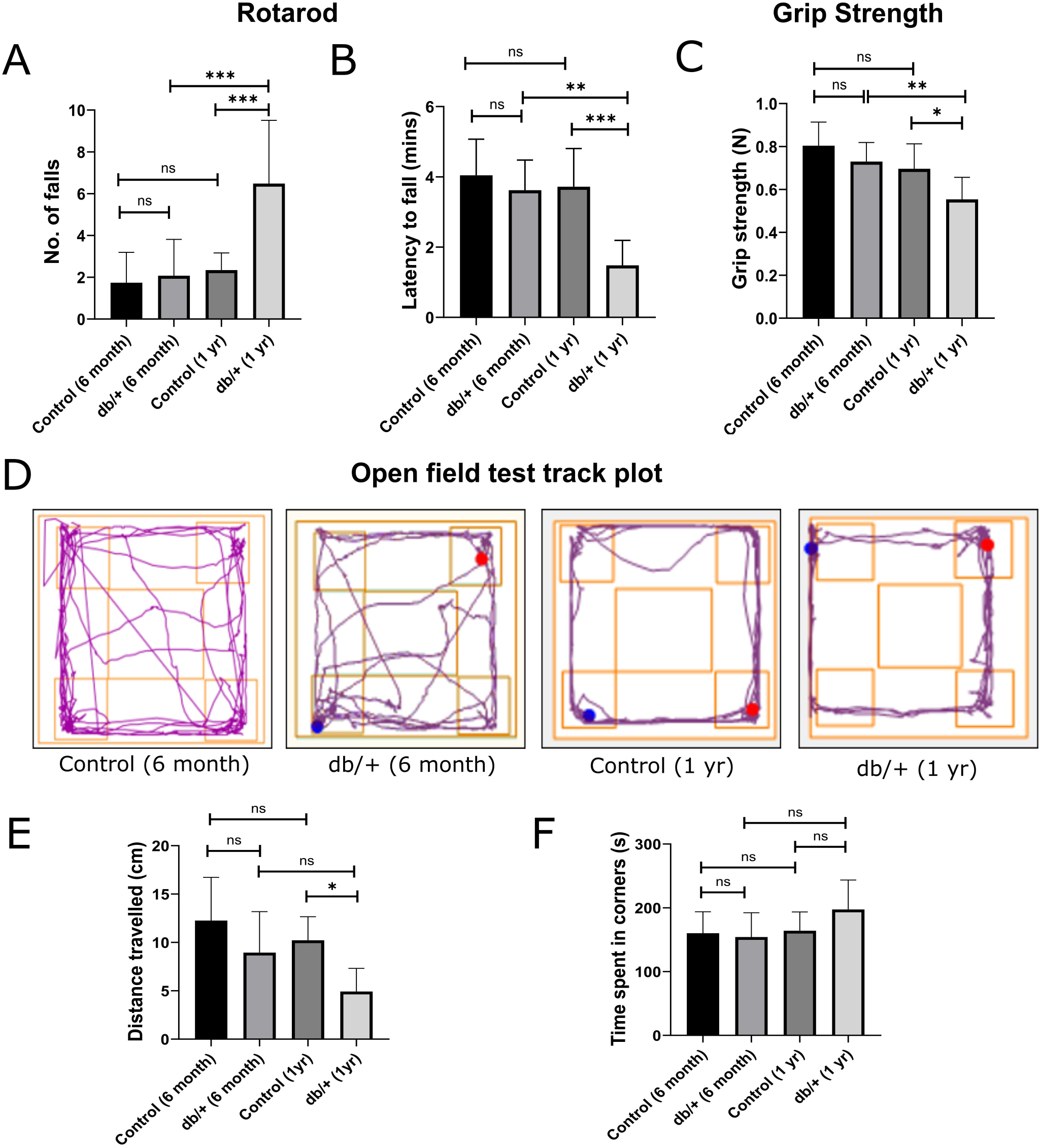
Aged db/+ mice show loss of muscle strength and decreased locomotor behavior. (A-B) Rotor-rod test for motor coordination: (A) Number of falls in duration of 5 minutes on the accelerated rotor rod, (B) time standing on the rotor-rod. (C) Grip strength was measured in Newtons, and there was an average of three trials per mouse. (D-F) Open field test for exploratory and locomotory behavior: (D) Representative track plot depicting the exploratory path of mice in an open field for the duration of five minutes. (E) The bar graph shows the distance traveled by mice in the open field. (F) The bar graph shows the time spent in the corners of the open field. All the groups have n=9 mice, and the data were presented as mean ± SD. The analysis was done using one-way ANOVA, and significance was denoted as *p>0.05, **p>0.01, and ***p>0.001.

To evaluate locomotor and exploratory behaviors, we conducted the open field test, which showed that one-year-old db/+ mice explored less, as indicated by a shorter total travel distance relative to control mice. This diminished exploratory drive is consistent with previous findings reporting depression-like traits in both diabetic patients and db/db mice (Figure 2D-F). Together, these observations suggest that the heterozygous Lepr mutation contributes not only to metabolic imbalance but also to accelerated age-related declines in motor and behavioral functions, thereby impacting overall physical and psychological well-being in aged mice.

### 4.3 Accelerated Cognitive Impairment in db/+ Mice: Short-Term and Long-Term Memory Deficits

To further explore cognitive impairments associated with the heterozygous *Lepr* mutation in db/+ mice, we employed the Morris water maze (MWM) and Y-maze tests to assess spatial learning and memory function. The MWM test offers a reliable measure of cognitive performance, as the drive to escape water is relatively independent of body weight or activity level. Moreover, swim paths in the MWM are not influenced by body weight or land-based motor performance, making it a reliable cognitive test.

At six months of age, db/+ mice performed comparably to wild-type controls in the MWM, with no significant differences in latency to reach the platform zone on the retrieval day, suggesting intact long-term memory at this stage. However, by one year of age, db/+ mice demonstrated a significant increase in latency to reach the platform zone, indicating a decline in long-term memory relative to age-matched wild-type mice (Figure 3A-B). To assess working memory, we employed the Y-maze test. At six months, both genotypes exhibited comparable spontaneous alternation behavior, indicating preserved short-term memory. In contrast, one-year-old db/+ mice showed altered exploration patterns, such as reduced arm entries or decreased time in specific arms, suggesting early signs of short-term memory impairment. Notably, spontaneous alternation percentages remained unchanged, implying that certain components of working memory may still be functional (Figure 3C).

**Figure 3.**
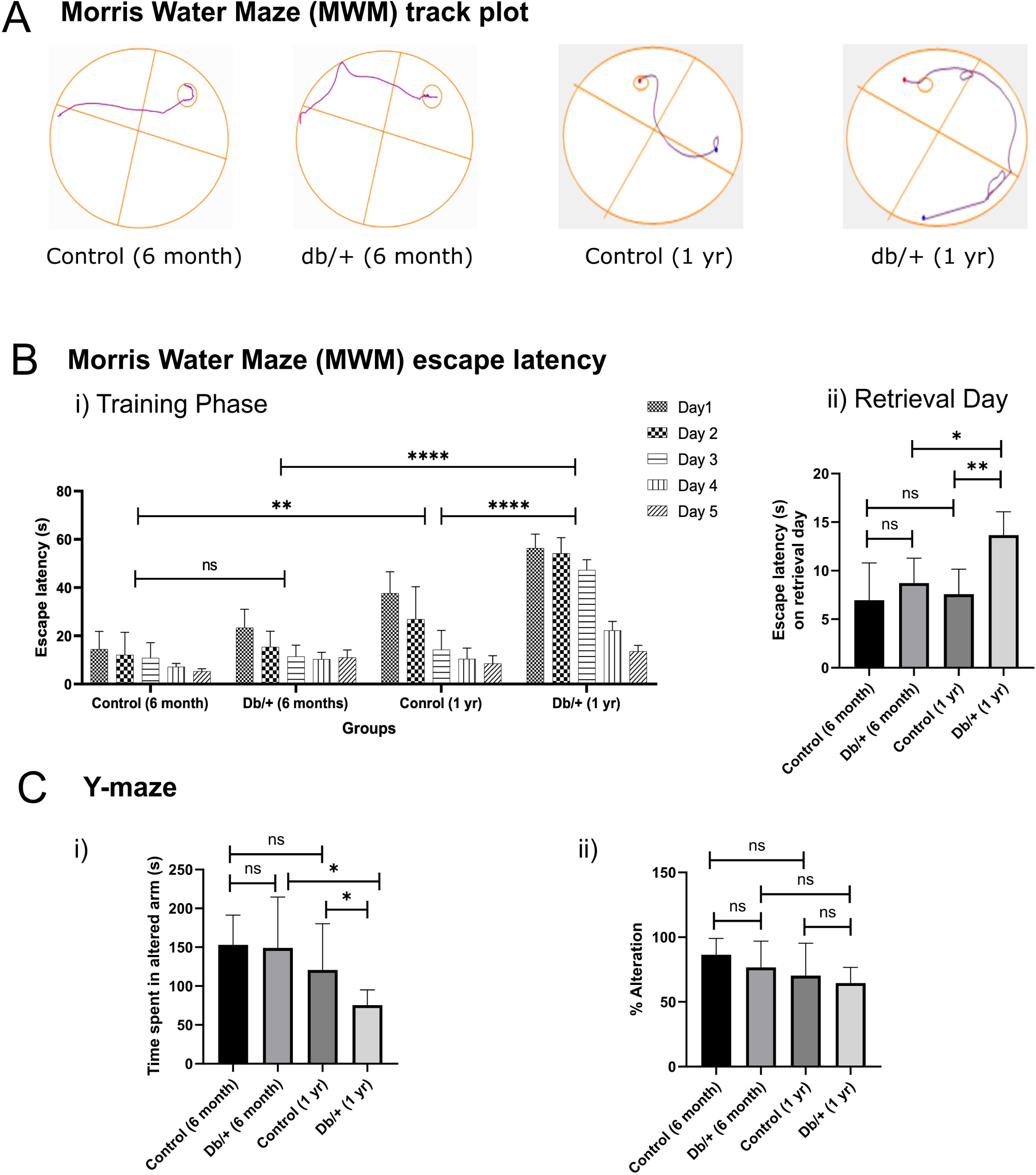
Short-term and long-term memory impairment is accelerated in aged db/+ mice. (A) Representative track plot of the MWM (day 5) of each group. (B (i)) Spatial acquisition in the Morris water maze, where the latency to reach the platform was scored over 5 days, wherein each day the mice received four trials. (B (ii)) Escape latencies on the retrieval day in seconds in MWM. (C) Y-maze to analyze short-term spatial working memory. (C (i)) Time spent in the altered arm in seconds. (C (ii)) Percentage alterations between the two arms. Values were presented as mean ± SD; n = 9, *p < 0.05, **p < 0.01, ***p < 0.001, and analysis was done using one-way ANOVA using GraphPad Prism.

These findings are consistent with earlier studies showing that diabetic models often display cognitive deficits. The emergence of both short- and long-term memory impairments in aging db/+ mice underscores the broader neurological consequences of the *Lepr* mutation. Beyond its established role in metabolic and physical decline, the mutation also appears to accelerate cognitive aging, highlighting its impact on brain function over time.

### 4.4 Aged Db/+ mice show elevated AD pathology and CDK5 upregulation

To investigate the molecular mechanisms underlying cognitive deficits in db/+ mice, we examined AD related markers in the brain. Pratap et al. ^17^ reported that in the AD mice model, Leptin signalling is dysregulated and is accompanied by plaque loading.

We first measured LEPR expression levels and found no significant differences between groups (Figure 4A). To assess whether LEPR signaling was altered in aged db/+ mice, we examined the phosphorylation of STAT3 at tyrosine 705 (pSTAT3-Y705) by western blot. A significant reduction in pSTAT3 levels was observed in aged db/+ mice, indicating impaired downstream signaling (Figure 4B).

**Figure 4.**
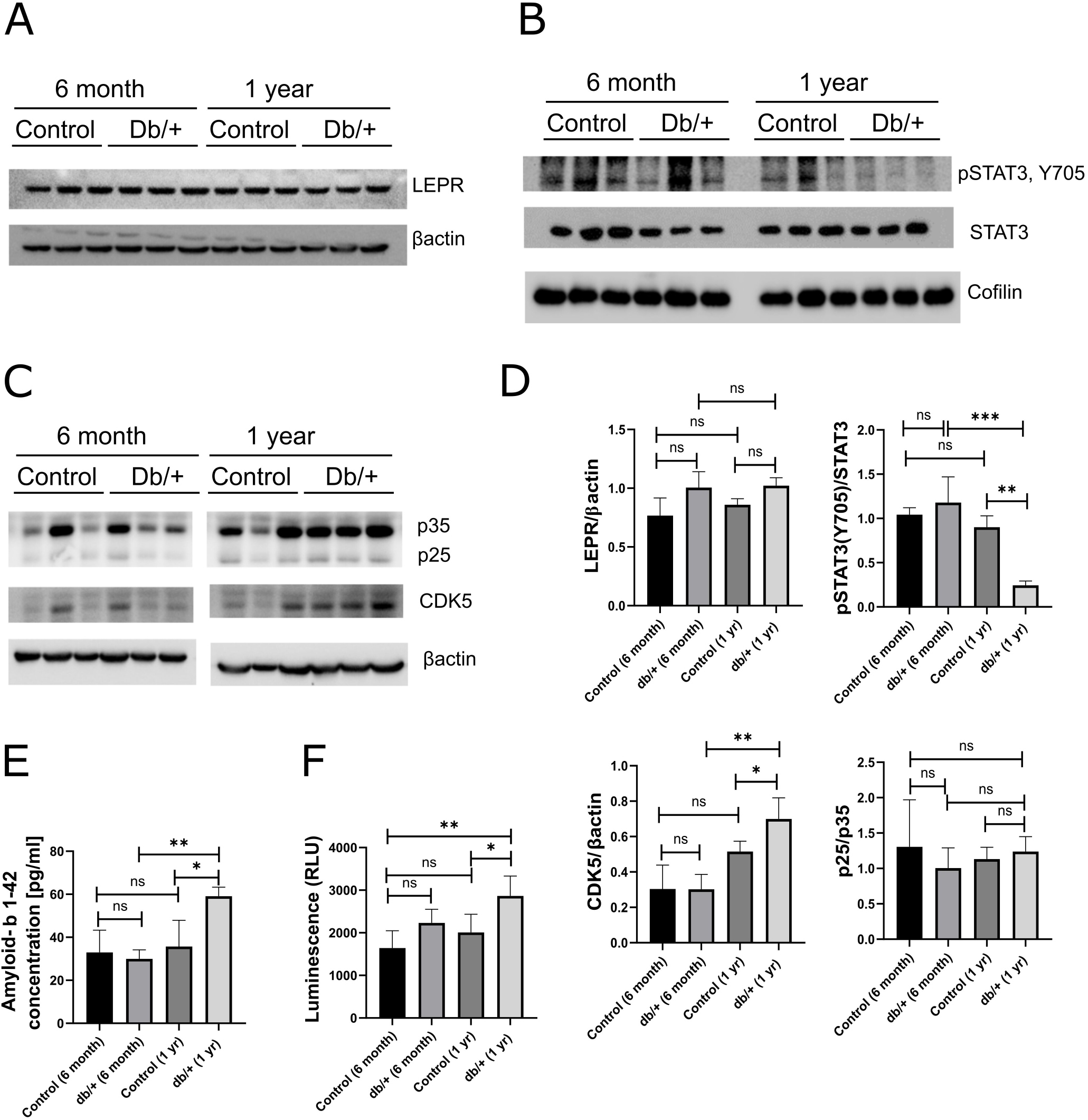
Aged db/+ mice show impaired STAT3 signalling and CDK5 upregulation. (A-D) Western blots of LEPR, STAT3, phosphorylated STAT3, p25, p35, and CDK5, and their corresponding densitometry normalised to their respective housekeeping protein. The expression of pSTAT3 and p25 was normalized to total STAT3 and p35, respectively. (E) The bar graph shows the ELISA quantification of Ab42 levels. (F) The bar graph shows CDK5 activity that correlates with the luminescence (in relative light units) with histone as a substrate of CDK5. All data are indicated as mean ± SEM and n=3. **p < 0.005 and ***p < 0.0005 determined by one-way ANOVA using GraphPad.

Next, we measured amyloid-beta levels using an enzyme-linked immunosorbent assay (ELISA), which revealed significantly elevated amyloid-beta in the brains of aged db/+ mice compared to age-matched controls (Figure 4E). Immunohistochemistry further confirmed increased Aβ42 accumulation in aged db/+ mice relative to both aged wild-type and young db/+ mice (Figure 5F).

**Figure 5.**
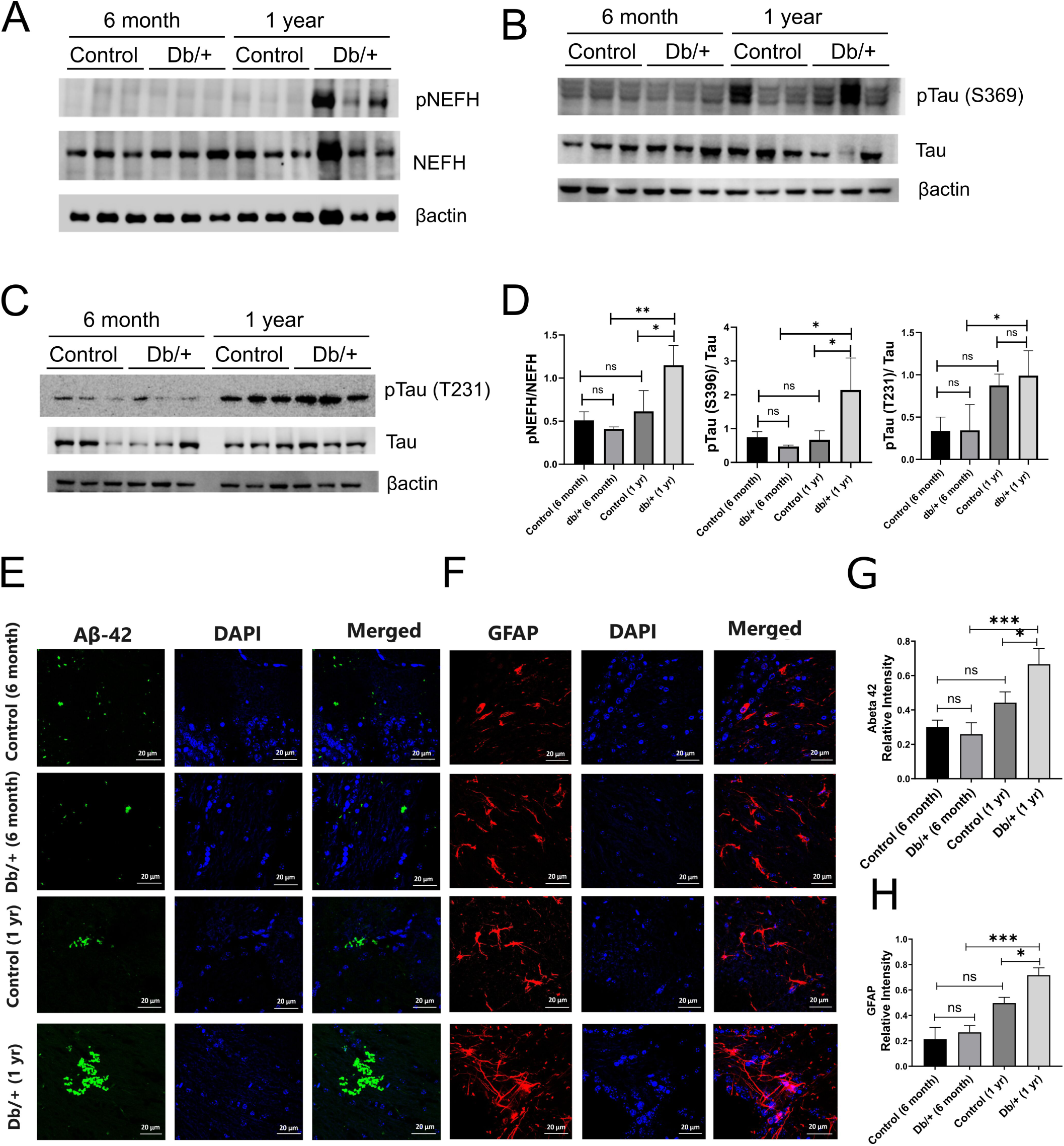
db/+ mutation accelerates tauopathy, Aβ-42 accumulation, and microglia activation in aged mice. (A-C) Western blots of NEF-H, pNEF-H, tau, and p-tau, and their corresponding densitometry, normalized to their respective housekeeping protein. The relative expression of phospho-NF-H and phospho-tau was normalized with the expression of total NF-H and total tau, respectively (therefore, the Y-axis value represents the ratio of protein expression of phospho-tau and phospho-NF-H to the protein expression of total tau and total NF-H). n=3 mice were used for western blot analysis. Immunohistochemistry of brain section (10um thickness) (D) Expressions of AB-42 (E) GFAP in the hippocampal region of the mouse groups. n=3 tissue sections were considered per group, along with n=3 images per section. (F)Relative intensity of Ab-42 and (G) GFAP with respect to DAPIs per group. The test used for the analysis was one-way ANOVA using GraphPad with data sets presented as mean ± SD. The significance were denoted as *p>0.05, **p>0.01; ***p>0.001.

Cyclin-dependent kinase 5 (CDK5), an atypical kinase, is found to be hyperactive in neurodegenerative disorders such as Alzheimer’s, Parkinson’s, and Huntington’s disease. CDK5 is also upregulated in the presence of amyloid plaques^18^. To assess whether amyloid beta in the brain of old db/+ mice causes a change in CDK5 activity and expression, we pulled down CDK5 from the brain lysates through immunoprecipitation and performed an *in vitro* kinase assay using histone as the substrate. Kinase assay of CDK5 revealed significantly high kinase activity in 1-year-old db/+ mice brains compared to the other groups (Figure 4F). We also checked p25 generation and CDK5 protein expression levels with western blot and observed the significant CDK5 upregulation in the brain of 1-year-old db/+ mice.

### 4.5 Db/+ mutation accelerates tauopathy and microglia activation in aged mice

Following the observed increase in CDK5 activity in aged db/+ mouse brains, we next investigated downstream pathological changes associated with CDK5 dysregulation. Since CDK5 is known to phosphorylate tau at specific residues implicated in tauopathy, we assessed tau phosphorylation at Ser396 and Thr231 by western blot. Aged db/+ mice exhibited significantly elevated levels of p-Tau (Ser396) and p-Tau (Thr231) compared to both age-matched wild-type and young db/+ mice, indicating enhanced tau pathology (Figure 5B and 5C).

To determine whether these changes extended to broader cytoskeletal alterations, we examined the phosphorylation status of neurofilament proteins. Western blot analysis revealed a marked increase in phosphorylated neurofilament heavy chain (pNF-H) in aged db/+ mice (Figure 5A). These findings suggest that the Lepr mutation contributes to both tau-based and neurofilament-based cytoskeletal disruption during aging.

In addition, we could see the activation of astrocytes, which correlates with enhanced GFAP expressions confirmed by immunohistochemistry studies in the brains of old db/+ mice compared with an age-matched control group and young mouse groups (Figure 5E and 5G).

### 4.6 Transcription profiling of old Db/+ mice

Next, bulk RNA-seq analysis was performed on brain tissues from 12-month-old db/+ mice to investigate regulatory genes involved in the progression of age-related cognitive impairment. Differentially expressed genes (DEGs) were identified using volcano plots, with thresholds set at p-value < 0.05 and absolute log₂ fold change (FC) > 2. In total, 1,496 genes were found to be differentially regulated—741 upregulated and 755 downregulated (Figure 6).

**Figure 6.**
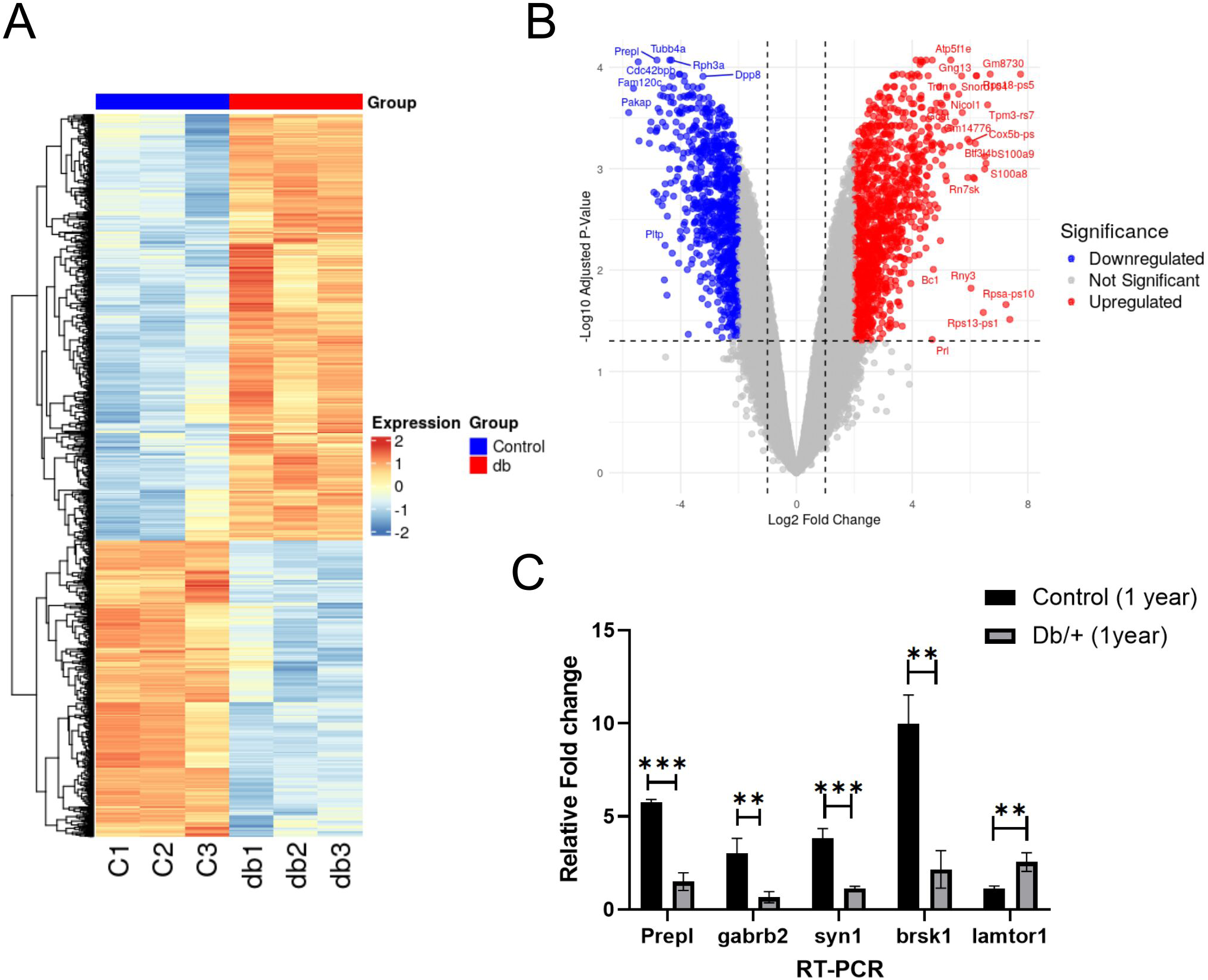
Transcriptome analysis of db mice and control mice. Total DEGs count is 1496, and Up:741 and Down: 755. (A) Heatmap of differential gene expression between the db and control mouse groups. (B) Volcano plots for the differentially expressed genes (DEGs). db vs control mice. (C) Measurement of mRNAs by RT-PCR. Five mRNAs (prepl, gabrb2, syn1, brsk1, and lamtor1) were randomly selected among those in (A) to confirm their expression change. P-value was calculated by a two-tailed t-test (n = 3 in each group).

Pathway enrichment analysis using the KEGG database revealed significant downregulation of pathways related to “neurotransmitter secretion,” “synaptic signaling,” and “learning and memory” in the cortex of db/+ mice compared to controls (Figure 7). Manual examination of these pathways showed marked reductions in the expression of genes such as Gabrb2, Brsk1, Camk2a, Syn1, and Syngap1 (Figure 6C). Reduced expression of these genes is associated with impaired memory due to disruptions in synaptic vesicle trafficking, neurotransmitter release, and AMPA receptor regulation—ultimately leading to weakened synaptic plasticity, altered excitatory-inhibitory balance, and impaired long-term potentiation (LTP). These findings were further validated by quantitative real-time reverse-transcription PCR (qRT-PCR).

**Figure 7.**
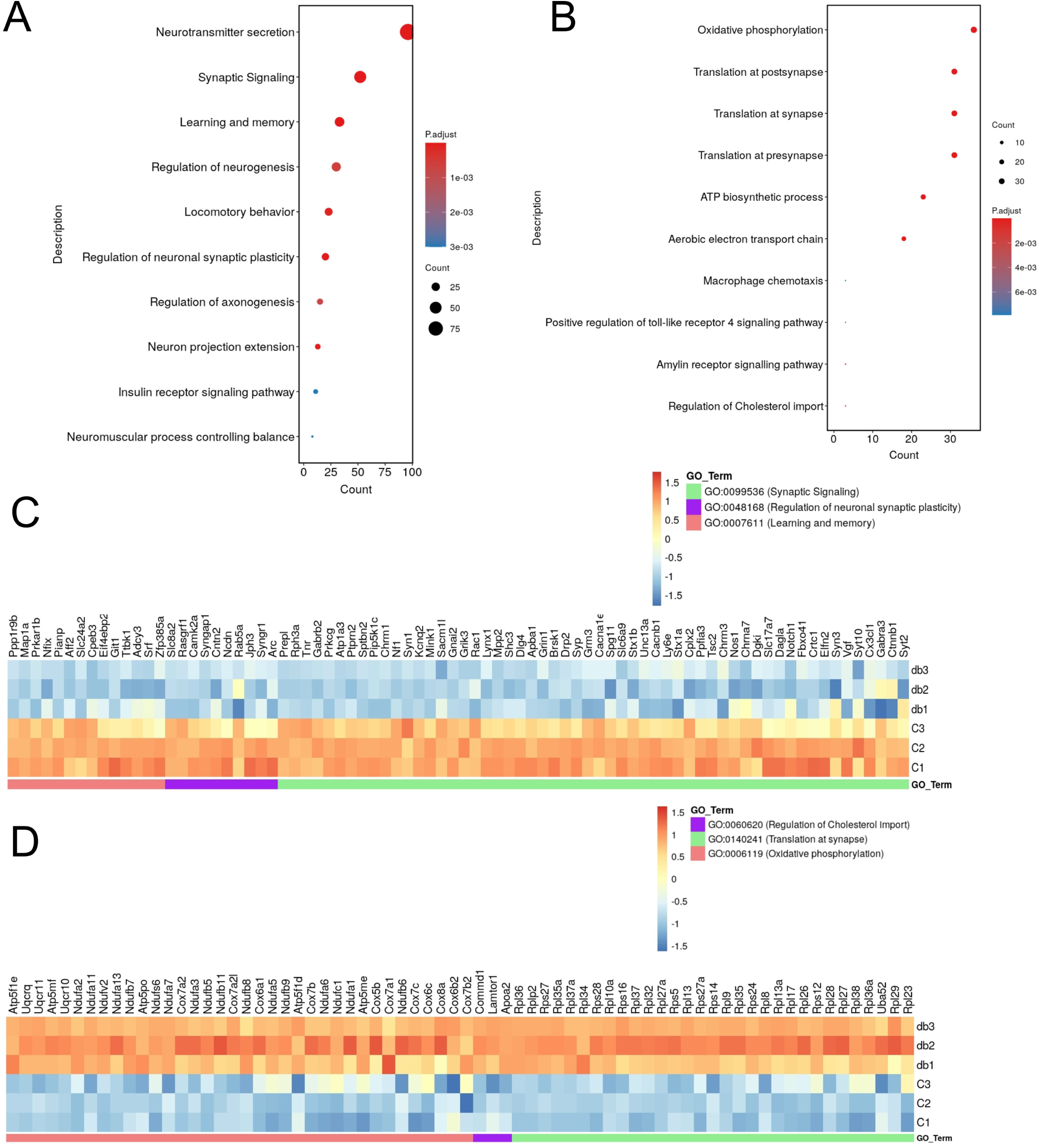
Heatmap for selected GO terms. (A) Dot plot of enriched GO terms for downregulated genes. (B) Dot plot of enriched GO terms for upregulated genes. (C) Heatmap depiction of the downregulated genes associated with GO:0099536 (Synaptic Signaling), GO:0048168 (Regulation of Neuronal Synaptic Plasticity), and GO:0007611 (Learning and Memory). (D) Heatmap depiction of the upregulated genes associated with GO:0060620 (Regulation of Cholesterol Import), GO:0140241 (Translation at Synapse), and GO:0006119 (Oxidative Phosphorylation).

Among the top upregulated pathways were “oxidative phosphorylation (OXPHOS)” and “translation at the pre- and post-synapse” (Figure 7). Overactive OXPHOS may disturb the energy homeostasis required for synaptic plasticity, thereby impairing processes such as LTP and long-term depression (LTD), which are essential for memory formation and maintenance.

### 4.7 *LEPR* variant curation and classification according to ACMG

To extend the relevance of our findings from the db/+ mouse model to human populations, we examined the genetic landscape of the human LEPR gene. Given that Lepr haploinsufficiency in mice led to age-related cognitive decline and Alzheimer-like pathology, we hypothesized that rare or under-characterized LEPR variants in humans might similarly contribute to neurodegenerative risk, especially in the context of metabolic disorders. Therefore, we systematically curated and analyzed LEPR variants from publicly available datasets and literature to evaluate their potential clinical significance. We compiled a total of 551 genetic variants of the LEPR gene from publications and databases associated with obesity and diabetes (Supplementary). These unique variants were categorized based on the type of mutation as follows: substitutions (357), insertions (11), deletions (10), and duplications (11). Further classification based on the predicted consequences of these variants at the gene and protein levels revealed multiple categories, including nonsynonymous, synonymous, UTR5, UTR3, upstream, stop gain, stop loss, splice site, frameshift deletions, non-frameshift deletions, and frameshift insertions, as summarised in (Figure 8D). All variants were systematically re-classified according to the ACMG & AMP standard guidelines. Of the 551 curated variants, 29 were classified as pathogenic, 6 as likely pathogenic, 111 as benign, 73 as likely benign, and 332 as variants of uncertain significance (VUS) (Figure 8C). A well-annotated and curated resource of LEPR genetic variants, such as this, may also facilitate a deeper understanding of population-specific patterns in obesity-associated genetic risk. On plotting allele frequency of pathogenic/likely pathogenic variants, we found that variant NM_002303.6:c.946C>A was highly prevalent in Middle Eastern and South Asian populations.

**Figure 8.**
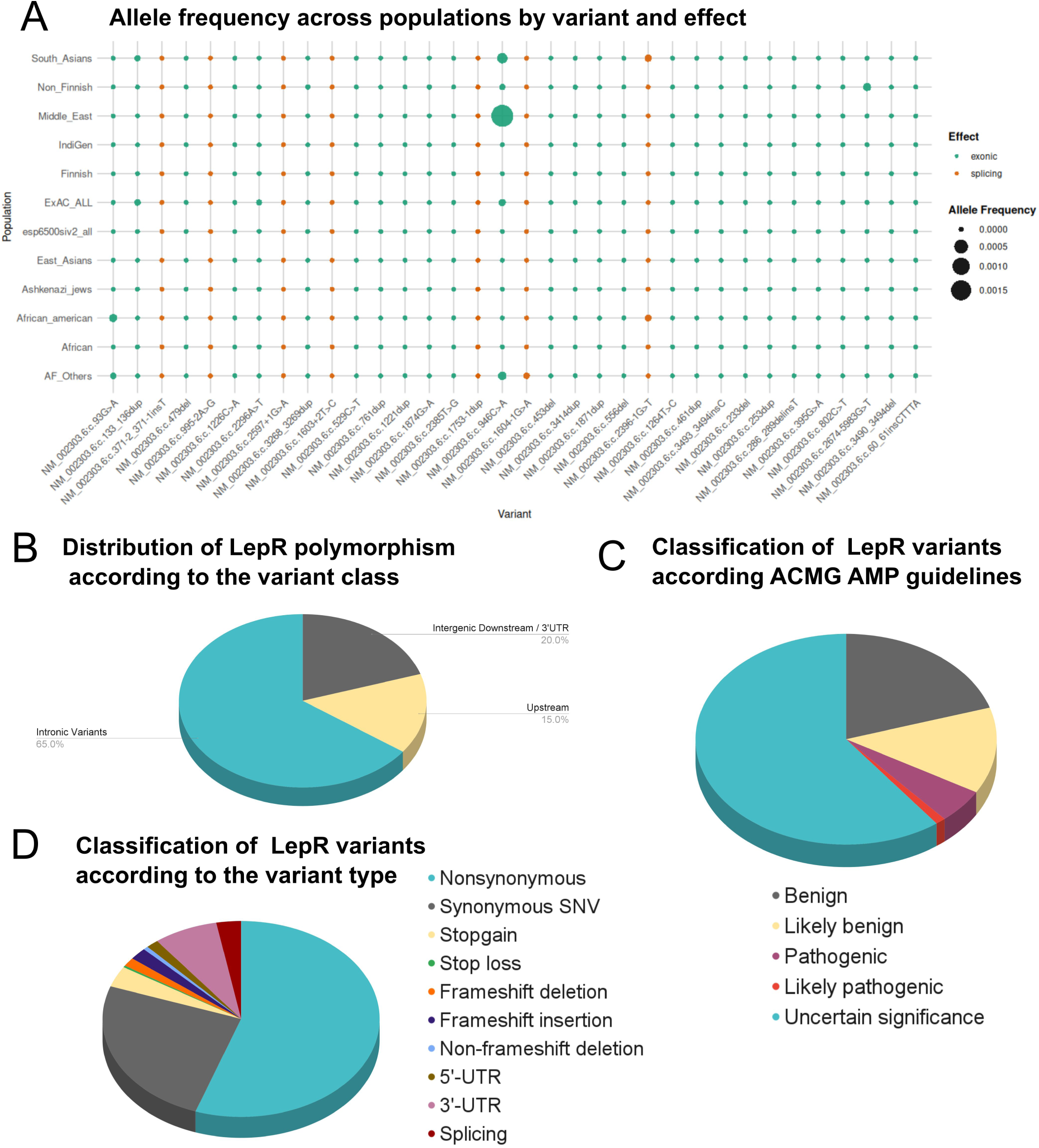
LEPR variant curation and classification according to ACMG. (A) Allele frequencies of the Pathogenic /Likely Pathogenic variants were queried across the dataset of population genomes and exomes from across the world (B) Distribution of LEPR polymorphism according to the variant class (C) ACMG Classification as pathogenic, likely pathogenic, benign, likely benign and VUS (Variant of Uncertain Significance) (D) Classification based on variant type.

## 3. DISCUSSION

The increasing prevalence of obesity in today’s population is alarming. Obesity is connected to various issues like cognitive decline and dementia. However, our understanding of how obesity causes damage to the central nervous system is incomplete and fragmented. Both animal models and human cohort studies found a high risk of developing cognitive impairment in obese individuals^19^. While unraveling the exact cause-and-effect relationships in these intricate disorders is challenging, rodent models have offered valuable insights into shared mechanisms. Db/db mice are one such model that lacks the leptin receptor’s long isoform due to an insertional mutation. Db/db mice develop a higher body weight, blood glucose and insulin resistance within 3-5 months compared to heterozygous db/+ mice, which are used as a control. Recently, it was reported that there is also an age-dependent development of obese phenotype in the db/+ mice ^7^.

Our study showed cognitive impairment and Alzheimer’s pathology in db/+ mice in an age-dependent manner. First, we measured the body weights of old and young db/+ and wildtype mice and found that db/+ mice show elevated body weights in old age compared to the age-matched controls. Aged db/+ mice also had elevated blood glucose and higher insulin resistance compared to the other groups. This result was in accordance with previous studies on db/+ mice ^8^. Next, we tested db/+ mice for muscle strength and found that aged db/+ mice showed significant muscle weakening compared to the other groups. It has been reported that a decline in muscle strength precedes tauopathy in mouse models ^20^, and greater grip strength is associated with better cognitive functioning ^21^. On analysing long-term and short-term memory with MWM and Y-maze test, our findings indicate that heterozygous leptin receptor mutation accelerates cognitive decline at the 1-year mark, as evidenced by poor performance in cognitive tasks.

Leptin has an important role in memory and learning. Leptin receptors are highly expressed in the hippocampus and cortex, where they facilitate memory-boosting long-term potentiation (LTP) and adaptability-enhancing synaptic plasticity ^22^. Since the leptin receptor needs to form a dimer to conduct intracellular signalling, a lower quantity of intact long receptor isoform molecules could lead to attenuated signals. We checked for LEPR protein expression in the db/+ mice but found no change in the protein levels. This might be due to the use of whole-brain lysate for Western blots. Since the leptin receptor is expressed differentially in different brain regions, capturing a subtle decrease in one isoform in one region would be difficult using whole-brain lysates. So, we checked for its downstream molecule, STAT3, and found a significant decrease in the phosphorylation levels of STAT3 (Y705) in the whole brain lysates of aged db/+ mice. This decrease in pSTAT3 indicates a decreased LEPR signaling in the brain. Since Leptin plays a protective role and prevents amyloid buildup in the brain by downregulating the expression of β- and γ-secretase ^23–25^, we checked for amyloid ß42 levels by ELISA and IHC. Our results show increased Aβ levels in aged Db/+ mice compared to the other groups.

Oligomerised Aß induces neurotoxicity and causes an increase in intraneuronal Ca2+ levels ^26^. An increase in intraneuronal Calcium triggers a pathological cascade involving calpain activation and cleavage of p35 into p25 ^27^. p25 then hyperactivates CDK5, which is a known kinase involved in Alzheimer’s pathology^14,28^. Hyperactive CDK5 then phosphorylates tau, forming neurofibrillary tangles. To confirm this, we first checked CDK5 levels and CDK5 activity and found them to be significantly upregulated in aged db/+ mice. Next, we looked for tau hyperphosphorylation at sites t231 and s396, which are CDK5 phosphorylation sites^29^. We found that there is a significant increase in tau phosphorylation in aged db/+ mice. Hyperphosphorylation of tau is one of the major pathological hallmarks of AD and other related neurodegenerative diseases. In AD, astrocytes become activated in response to the amyloid plaques and tau tangles, the two hallmark pathologies of the disease. This activation process, known as reactive astrogliosis, involves increasing GFAP production. GFAP levels were also found to increase significantly in aged db/+ mice in our study. In contrast, 6-month-old db/+ mice showed no cognitive impairment, no increase in ab42, no CDK5 upregulation, or tauopathy.

Subsequently, RNA-seq of brain tissue revealed that Gabrb2, a key factor associated with Alzheimer’s disease (AD), was found to be downregulated in aged db/+ mice ^30^. Gabrb2 encodes the Gamma-Aminobutyric Acid Type A Receptor Subunit Beta2, a critical component of GABAergic signaling, which mediates the fastest inhibitory synaptic transmission in the mammalian brain^31^. Reduced gabrb2 leads to fewer functional GABA_A receptors, weakening inhibitory signaling and causing an imbalance between excitation and inhibition. Gene ontology (GO) analysis further supported this by revealing a downregulation of the neurotransmitter secretion pathway in the brains of db/+ mice. Additionally, brsk1 was also reported to be downregulated in the db/+ group. Brsk1 phosphorylates synaptic vesicle proteins and modulates AMPAR and NMDAR trafficking, which is critical for long-term potentiation ^32^. Brsk1 also regulates presynaptic vesicle docking and fusion by interacting with SNARE proteins, which ensures efficient glutamate and GABA release, balancing excitatory and inhibitory signaling, which is crucial for cognitive function. Several other important players in memory formation and synaptic plasticity, like camk2a, syn1, syngap,1 were also found to be downregulated in old db/+ mice. The neuronal network dysfunction hypothesis has recently rekindled interest in understanding how disruptions in neuronal connectivity at the network level result in epileptiform activity seen in the brains of AD patients and mouse models ^33^. Therefore, learning and memory-related neural network deficiencies may eventually lead to memory problems. In this case, it has been proposed that a mismatch between excitation and inhibition is a factor in the malfunctioning of neural networks. Other pathways that were differentially upregulated in db/+ mice were “oxidative phosphorylation (OXPHOS)”, “translation at pre- and post-synapse”, and “ATP biosynthesis process”. Upregulation of translation can overwhelm protein quality control mechanisms, leading to misfolded proteins and toxic aggregates that impair synaptic function. Additionally, overproduction of synaptic proteins disrupts their stoichiometric balance, resulting in aberrant signaling and impaired plasticity, while increased ATP demand exacerbates oxidative stress, further damaging synapses. These findings suggest that leptin receptor deficiency disrupts critical synaptic processes, potentially contributing to the cognitive and memory impairments observed in these mice.

To extend the relevance of our findings to humans, we curated all the leptin receptor variants from publicly available databases and published literature. Variants that were classified as likely pathogenic/VUS were reclassified as pathogenic. We further looked at the distribution of these pathogenic and likely pathogenic variants by curating allele frequency across different datasets. We found that the likely pathogenic variant NM_002303.6:c.946C>A was high in the Middle East and South Asian populations. Since *LEPR* mutation-associated obesity cases are rare, there can be two reasons for such a high prevalence of this particular mutation. First, genetic testing for obesity has not reached the mainstream in the Middle East and South Asian countries, and second, it can be present in heterozygous conditions. Thus, the phenotype does not manifest at an early age.

This detailed LEPR variant curation directly connects with the central theme of the study, the impact of heterozygous *Lepr* mutations on aging-related neurodegeneration and metabolic dysfunction and reinforces the translational relevance of the db/+ mouse model. In this model, aged db/+ mice with partial leptin receptor deficiency exhibited a range of Alzheimer’s disease (AD)-like pathologies, including cognitive deficits, CDK5 hyperactivation, tau hyperphosphorylation, and increased amyloid-β42 accumulation. These pathological hallmarks are strongly aligned with human AD pathology and are mechanistically linked to disrupted leptin signaling. Importantly, impaired phosphorylation of STAT3 (Y705) in aged db/+ brains—an indicator of defective LEPR downstream signaling—was observed despite unchanged total LEPR protein levels. This supports the notion that the functional integrity of LEPR signaling, rather than its expression alone, is critical for neuroprotection. The parallel identification of 33 pathogenic or likely pathogenic human *LEPR* variants, with several showing high allele frequencies in specific populations (e.g., South Asian and Middle Eastern), offers a compelling molecular rationale for stratifying at-risk individuals based on genotype. Furthermore, transcriptomic analysis of aged db/+ brains revealed significant downregulation of genes involved in synaptic signaling, neurotransmitter secretion, and learning and memory (e.g., *Gabrb2, Brsk1, Camk2a, Syn1, Syngap1*), alongside upregulation of oxidative phosphorylation (OXPHOS) and synaptic translation pathways. These molecular shifts point to a dysregulated synaptic environment, potentially exacerbated by metabolic stress and altered energy homeostasis. The combined effects of altered gene expression, neuroinflammation, CDK5/p25-driven tau pathology, and impaired LEPR-STAT3 signaling converge to form a pathophysiological axis that mirrors findings from the human variant analysis. Notably, the presence of variants of uncertain significance (VUS) in the *LEPR* gene suggests that there may be many underappreciated or context-dependent contributors to cognitive decline that manifest later in life, particularly in individuals exposed to metabolic challenges like obesity or type 2 diabetes. Thus, the convergence of mouse phenotype, mechanistic data, and human genetic variation strongly supports a model wherein partial LEPR loss constitutes a genetic vulnerability factor for age-related cognitive impairment and neurodegeneration. This emphasizes the need for integrative approaches combining functional genomics, phenotypic characterization, and population screening to uncover hidden genetic risks associated with common disorders like AD.

## 4. METHODS

### 2.1 Animals

All animal experiments were carried out in accordance with protocols that had been approved by the CSIR IGIB’s Institutional Animal Care and Use Committee. Mice were housed in groups of 5 to 6 per cage under standard humidity and temperature conditions, and a 12:12–hour light–dark cycle was maintained throughout the study. 4-5 weeks old male C57/BL (n=9) and db/+ (n=9) mice were divided into two groups. One group from both C57/BL and Db/+ was sacrificed when they were 6 months old, and another was sacrificed at the age of 1 year. Body weight, fasting blood glucose, glucose tolerance, and insulin tolerance were determined at 6 months and 1 year of age.

### 2.2 Behavioral Paradigms

All the behavioral paradigms were carried out in the dark phase of the housing conditions. The mice were transferred to the behavior testing room an hour before the testing began to habituate the animals to the testing room. The apparatus was cleaned with 70% alcohol after every individual session, such that there were no remains of traces or odour of the previously tested animal.

### 2.3 Morris water maze (MWM) test

The test was performed as described [V Vorhees C, 2006] with modifications. A white 150 cm diameter polyethylene tank filled halfway with room temperature water (21°C ± 1°C) was used. A white platform was submerged 1.5 cm below the surface of the water, and the size of the platform varied depending on the test phase. Mice were tested in rotation, with the inter-trial interval determined by the number of mice being tested at any given time; however, the time was approximately 10 min. Latency to reach the platform and path efficiency (straight-line distance from start to platform ÷ the path taken by the mouse) to reach the goal, along with mean swim speed, were measured using Any-Maze video tracking software (Stoelting Instruments, Wood Dale, IL).

### 2.4 Grip strength

Grip strength is a non-invasive method to test muscle strength in rodents [O.A. Meyer, H.A. Tilson, Neurobehav. Toxicol. 1 (1979) 233–236. The forelimb grip strength test was performed using a grip strength meter. The mouse was allowed to grasp the gauge with its forelimbs, and then its tail was slowly pulled backwards. The muscular force was recorded on a digital transducer. Each mouse was subjected to 5 trials.

### 2.5 Open field test

The open field chamber consisted of a white square arena (60*60cm). The experimental procedure was carried out in a well-lit, centrally placed overhead red light. The test was carried out in 2 consecutive days - day 1-habituation to the arena, and Day 2-testing phase. The habituation phase lasted for 5 minutes for each animal, and the testing phase on Day 2 lasted for 10 minutes each. The animals were acclimatized to the room an hour before the testing commenced. Animals were placed in the center of the arena, and their exploratory behavior was recorded through Anymaze software [V. Carola, F. D’Olimpio, Behav. Brain Res. 134 (2002) 49– 57.]

### 2.6 Y-maze test

The Y-maze test was used to evaluate spatial working memory based on spontaneous alternation behavior. The apparatus consists of three identical arms arranged at 120° angles. Mice were tested during their active dark phase in a dimly lit, undisturbed room. Each animal was allowed to freely explore the maze for 5 minutes, and their behavior was recorded using Anymaze tracking software. An alternation was defined as consecutive entries into all three arms without repetition. Healthy mice are expected to show a higher rate of alternations, reflecting an ability to remember previously visited arms and a natural tendency to explore novel environments ^12^.

### 2.7 Rotarod Test

Rotarod test is used to assess Motor coordination in mice. The animal is placed on a horizontal rod that rotates about its long axis, and the animal must walk forward to remain upright and not fall off.

### 2.8 Immunoprecipitation

CDK5 was immunoprecipitated from mouse brain lysate. CDK5 monoclonal antibody (Invitrogen-) was incubated with Agarose A/G beads for 6 hours at 4°C, followed by washing with lysis buffer. 200 μg of brain lysate was added to the beads and incubated overnight at 4°C, and then proceeded with washing with a lysis buffer to remove the unbound protein. The elution was done in 0.2 M glycine buffer.

### 2.9 Kinase Assay

Determination of CDK5 kinase activity was performed using the Promega ADP-GloTM Kinase Assay kit. It is a luminescence-based assay, where the kinase activity is measured by detecting the amount of ADP produced during the kinase reaction. Initial experiments were carried out to check the activity of Cdk5/p35. To serve this purpose, 2ug of histone was used as a substrate for the enzyme complex. 100 uM ATP was added to each reaction. The luminescence generated from the reaction is directly proportional to the amount of ADP produced and, consequently, the kinase activity ^4,13–15^.

### 2.10 Glucose Tolerance test and Insulin Tolerance test

2g/kg glucose for glucose tolerance test, and 0.75U/kg human insulin was used for insulin tolerance test. The glucose levels were measured using a one-touch glucometer at different time points (30, 60, and 120 mins) after the administration of glucose and insulin ^16^.

### 2.11 Immunohistochemistry

Brain tissues preserved in 4% formaldehyde were processed for paraffin embedding. Thin sections of 5 µm were cut from the paraffin blocks and mounted onto adhesive-coated glass slides. Immunohistochemistry was performed following established protocols for single and double staining. The primary antibodies used included Aβ 1-42 (1:100) and GFAP (1:300). Fluorescent signals were visualized using Alexa Fluor-conjugated secondary antibodies (Invitrogen), which emit at 488 nm (green) and 594 nm (red) ^4,13^.

### 2.12 Western Blotting

Mice were sacrificed according to the Institution’s ethical procedures. Protein lysates from the mouse cortex were homogenized in ice-cold T-per lysis buffer containing protease inhibitor cocktail and phosphatase inhibitor cocktail, and the amount of protein in lysates was estimated using BCA reagent (Thermo Fisher Scientific, Rockford, IL, USA). Protein samples were separated using SDS-PAGE and then transferred to a nitrocellulose membrane. The membrane was then probed with antibodies as described previously. The primary antibodies were diluted in 1X PBS with 0.1% (v/v) Tween 20. (Antibodies used are listed in figure-). All electrophoresis and immunoblotting reagents were purchased from Bio-Rad Laboratories (Hercules, CA, USA).

### 2.13 Aβ analysis

The amount of Amyloid beta 1-42 was determined using ELISA KIT (Catalog no.-E-EL-M3010). One hemisphere of the mouse brain was homogenized and used in this experiment. ELISA data was analyzed using GainData (Arigo Biolaboratories).

### 2.14 RNA isolation

Total RNA was isolated from tissues preserved in RNAlater (Invitrogen) using TRIzol reagent (Invitrogen) following the manufacturer’s protocol. Tissues were stored in RNAlater at –80°C until processing. For RNA extraction, approximately 50–100 mg of tissue was removed from RNAlater, briefly blotted to remove excess solution, and homogenized in 1 mL of TRIzol using a mechanical homogenizer. The homogenate was incubated at room temperature for 5 minutes to permit complete dissociation of nucleoprotein complexes. Phase separation was achieved by adding 0.2 mL of chloroform per 1 mL of TRIzol, followed by vigorous shaking for 15 seconds and incubation at room temperature for 2–3 minutes. Samples were centrifuged at 12,000 × g for 15 minutes at 4°C to separate the phases. The upper aqueous phase containing RNA was carefully transferred to a fresh RNase-free tube, and RNA was precipitated by adding an equal volume of isopropanol. After incubating at room temperature for 10 minutes, the samples were centrifuged at 12,000 × g for 10 minutes at 4°C. The RNA pellet was washed with 1 mL of 75% ethanol, vortexed briefly, and centrifuged at 7,500 × g for 5 minutes at 4°C. The pellet was air-dried for 5–10 minutes and resuspended in RNase-free water. RNA concentration and purity were assessed using a NanoDrop spectrophotometer (Thermo Fisher Scientific), and integrity was confirmed via 1% agarose gel electrophoresis.

### 2.15 Transcriptome Analysis

RNA-Seq data analysis was performed following standard protocols. The reference genome for Mus musculus (GRCm38.dna.fa) was obtained from the Ensembl database (http://www.ensembl.org). Raw sequencing reads were pre-processed using Fastp (v0.23.2) to remove low-quality reads and adapters, and the quality of the filtered reads was assessed with MultiQC. A reference genome index was generated using HISAT2, and the high-quality reads were aligned to the reference genome with the same tool. Gene-level read counts were quantified using FeatureCounts, producing the raw gene expression matrix.

Differential expression analysis was performed using the edgeR package in R. Raw counts were normalized and the negative binomial statistical model implemented in edgeR was applied to identify differentially expressed genes (DEGs). Genes with an adjusted p-value (FDR) < 0.05 and an absolute log2 fold change (|log2FC|) ≥ 2 were considered significantly altered. To gain insights into biological pathways and molecular functions, the resulting DEGs were subjected to functional enrichment analysis using Metascape (https://metascape.org/), including Gene Set Enrichment Analysis (GSEA) and pathway-level clustering.

### 2.16 Quantitative real-time PCR

Total RNA was extracted from 30 mg of mouse cortex using TRIzol™ Reagent (Thermo Fisher Scientific). The corresponding cDNAs were prepared from 2 μg of extracted total RNA with a High-Capacity cDNA Reverse Transcription Kit (Applied Biosystems™), according to the manufacturer’s instructions. For qPCR, aliquots of cDNA templates were subjected to qPCR using PowerUp™ SYBR™ Green Master Mix⁷ (Applied Biosystems™), according to the manufacturer’s instructions. Primers were ordered from Sigma-Aldrich. Each sample was loaded in triplicate, and negative and positive controls were included. 18s was amplified as an internal reference gene. The following PCR conditions were employed using QuantStudio™ 7 Flex Real-Time PCR System: 50 °C for 2 min, 95 °C for 0 min, and 40 cycles each with 95 °C for 15 s and 60 °C for 1 min. The ΔΔCt method was used to analyze the data. The relative mRNA expression of each gene is reported as the mean and standard deviation of 3 independent total RNA extractions and real-time PCR analyses.

### 2.17 Statistical Analysis

For all other data, GraphPad Prism 9 (GraphPad Software) was used for statistical analysis and graphical representations. The appropriate statistical tests were chosen on the basis of experimental design after consulting the GraphPad Statistics Guide. Sample size was determined through power analysis, indicating that 9 mice per group were sufficient to detect significant behavioral differences, based on an estimated effect size (Cohen’s d = 1.2) derived from preliminary data, with 80% statistical power and a significance level of 0.05. This sample size balances the need for detecting meaningful effects while minimizing animal use in accordance with ethical standards.

## 5. ACKNOWLEDGEMENT

We would like to thank Swaraj Ranjan Paul for his contribution to the initial curation of Lepr gene variants. The authors gratefully acknowledge financial support from the Council of Scientific and Industrial Research (CSIR), India through grant OLP002501, and from the Science and Engineering Research Board (SERB), India through grant GAP0168. We also acknowledge fellowship support: a Senior Research Fellowship to Sangita Paul and Juhi Bhardwaj from CSIR, and a Senior Research Fellowship to Srishti Sharma from UGC.

## 7. THE PAPER EXPLAINED

### PROBLEM

The study addresses the problem that, although leptin signaling is known to have neuroprotective effects and is implicated in Alzheimer’s disease (AD), most research has focused on homozygous mutations of the leptin receptor (Lepr). This leaves a major gap in understanding how partial loss of leptin signaling, such as in heterozygous mutations, might influence age-related neurodegeneration. Since many human populations are more likely to carry heterozygous rather than homozygous mutations, clarifying their impact is crucial for assessing genetic risk in AD.

### RESULTS

In this study, we evaluated one-year-old db/+ mice, which carry a heterozygous mutation in Lepr. We found that these mice displayed both metabolic and neurological impairments: increased body weight and insulin resistance, memory deficits, elevated Aβ_42_ levels, tau hyperphosphorylation, CDK5 hyperactivation, and astrocyte activation. Transcriptomic analyses further revealed dysregulation of synaptic and mitochondrial pathways, linking leptin signaling to neuronal health at the systems level. Complementing the mouse data, we also identified thirty-three human LEPR variants that were classified as pathogenic or likely pathogenic using ACMG guidelines.

### IMPACT

The impact of these findings is significant. They demonstrate that even partial loss of leptin signaling (haploinsufficiency) is sufficient to trigger age-related cognitive decline and AD-like pathology, broadening our understanding of metabolic contributions to neurodegeneration. By establishing LEPR haploinsufficiency as a genetic risk factor, the study highlights leptin signaling as a potential therapeutic target. This shifts the perspective from considering leptin primarily as a metabolic regulator to recognizing its central role in maintaining brain health and preventing neurodegenerative disease.

